# The discovery of an external bacterial niche reframes sequencing-based microbiomes in a freshwater sponge

**DOI:** 10.64898/2026.02.07.704605

**Authors:** Jacob S. Pawley, Scott A. Nichols

## Abstract

Animals maintain beneficial associations with microbes while avoiding parasitism, often by physically segregating microbes from host tissues. However, sponges (Porifera) are frequently cited as an exception, with bacteria being shown to reside directly within the host mesohyl—an interior space composed of extracellular matrix and migratory cells. Yet this model rests largely on studies of field-collected specimens, where experimental control and spatial resolution are limited. Here, we exploit the experimental advantages of the freshwater sponge *Ephydatia muelleri*—culturable in the laboratory and highly amenable to confocal imaging—to directly address the spatial organization and transmission of host-associated bacteria. Contrary to expectations from studies of marine sponge systems, and to predictions from microbiome sequencing studies of freshwater sponges, we found limited evidence for diverse bacterial communities inhabiting the mesohyl of *E. muelleri*. Instead, we identified a novel microbial niche that is external to sponge tissues—a host-secreted matrix that underlies the basal epithelium, coats spicules, and envelops gemmules. In both laboratory cultures and field-collected specimens, matrix-associated microbes were abundant but separated from the sponge interior (mesohyl) by a continuous epithelial barrier. These external matrix-associated communities may contribute to the molecular signatures of stable freshwater sponge microbiomes, even if their functional roles remain unresolved. These findings reframe sequencing-based reports of freshwater sponge microbiomes with confocal imaging, and highlight that spatial context is essential for interpreting animal host–microbe associations.

Graphical Abstract
Microbial communities colonize the skeletal matrix during dormancy in *E. muelleri*.
In contrast to interpretations from microbiome sequencing studies, we find limited evidence for a resident microbiome within the mesohyl (shown in green) of the freshwater sponge, *Ephydatia muelleri*. Instead, we find that microbes are primarily confined to a novel external niche—a host-secreted matrix (shown in brown), which is exposed to the environment during periods of dormancy, and therefore permissive to microbial colonization. This matrix confers attachment to the basal substrate, embeds siliceous spicules, and coats gemmules (stress-resistant propagules maintaining senesced tissues). We find no evidence of bacteria being maintained across dormancy through the interior of gemmules. Instead, the external matrix and associated microbes are physically separated from the host mesohyl by a continuous epithelium (depicted as thick black lines). With spatial context afforded by in vivo imaging, we propose that these external-matrix-resident microbes likely account for previous reports of a stable, internal microbiome in freshwater sponges.

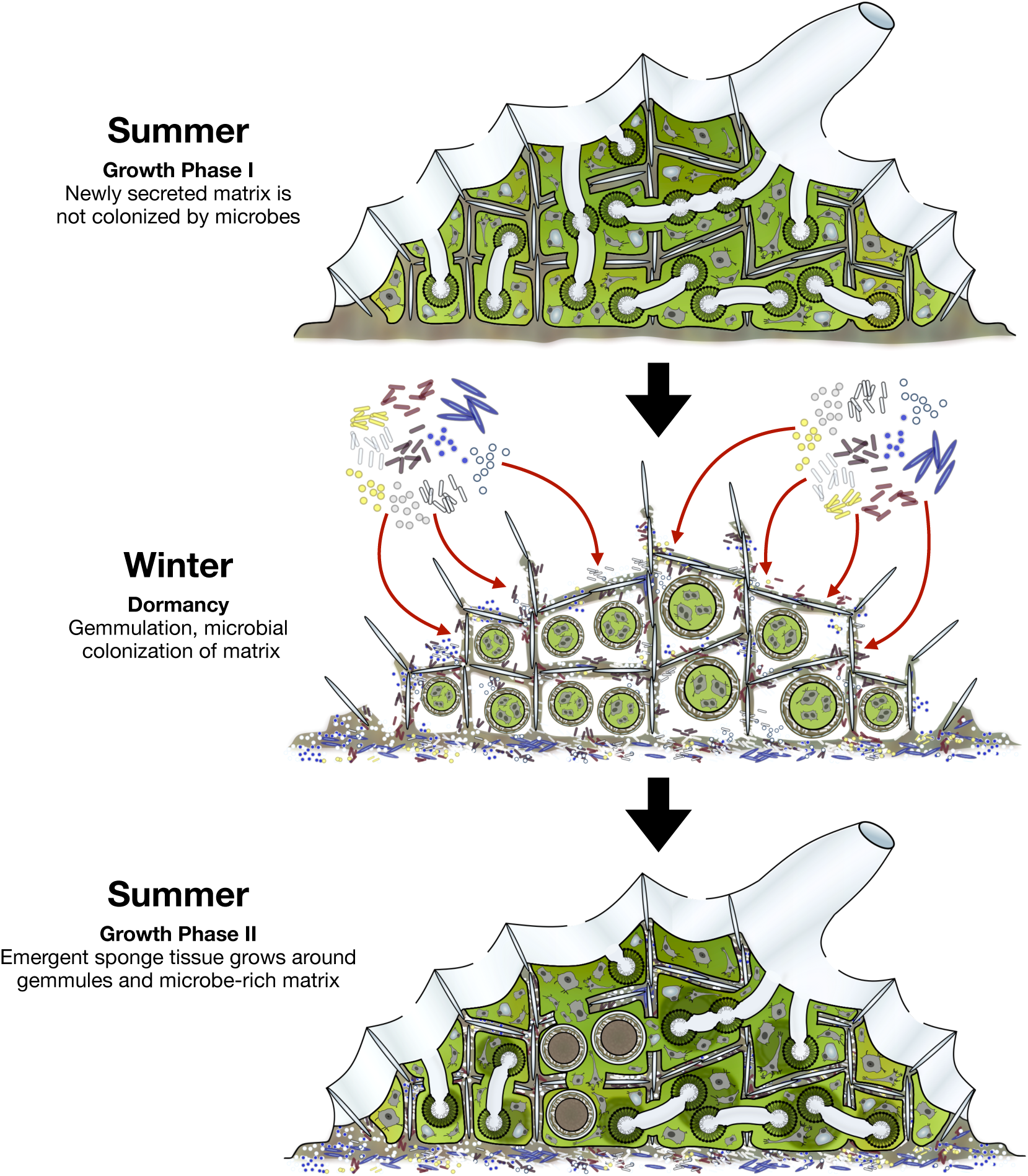

## Introduction

Animals often form stable, beneficial associations with microbes that are essential for normal physiology and development [1–3]. These relationships, however, exist under persistent tension: the host must accommodate cooperative symbionts while also defending against parasitism. A central mechanism that enables this balance is physical segregation, whereby microbes are excluded from the host body interior and instead confined to external surfaces, luminal spaces bounded by mucosal epithelia, or intracellular compartments derived from the host endomembrane system (e.g., symbiosomes in corals) [1, 4–7]. In general, when microbes breach these spatial constraints they are perceived as parasites and targeted by the host immune system. Stable symbioses are therefore confined to symbiont-permissive compartments that are spatially segregated from host-privileged tissues, allowing immune surveillance to protect the host interior without disrupting beneficial partnerships.

Sponges (phylum: Porifera) represent a striking departure from this organizational principle. In many marine species, dense and diverse microbial communities reside directly within the mesohyl—the internal compartment of the sponge body composed of extracellular matrix, interspersed with migratory host cells, and bounded by epithelia [8–11]. However, electron microscopy has revealed considerable variation in this pattern, with the mesohyl of some sponges being densely inhabited by bacteria and other species having very few, giving rise to the widely adopted high-microbial-abundance (HMA) versus low-microbial-abundance (LMA) framework [8, 12–16]. HMA sponges typically contain 10⁸–10¹⁰ microbial cells g⁻¹ and maintain phylogenetically complex, host-enriched consortia that include sponge-associated lineages such as *Poribacteria* and *Tectomicrobia* [17–21]. In contrast, LMA sponges harbor 2–4 orders of magnitude fewer microbes, levels similar to that of surrounding seawater (∼10⁵–10⁶ microbial cells g⁻¹), which are often enriched with transient or environmentally acquired taxa [22, 23]. Within Porifera, the extent to which a host permits microbial occupation of host-privileged compartments represents a trade-off linked to energy balance and filtration architecture [14].

A central limitation for advancing sponge microbiome research toward mechanistic understanding is that most species are difficult to cultivate under controlled laboratory conditions, heavily restricting functional studies [24]. Notable exceptions include freshwater sponges (e.g., *Ephydatia* and *Spongilla*), which regenerate from gemmules—stress-resistant, dormant propagules that can be hatched under fully controlled laboratory conditions [25–27]. Composed of >500 stem cells encased in a layered coat of collagen-like fibers and siliceous spicules, gemmules are anchored to the residual skeletal matrix from senesced tissues and remain viable until environmental conditions improve [28–30]. In culture, individual gemmules can regenerate into 1–2 mm sponges with transparent tissues, suitable for high-resolution imaging and experimental manipulation [24, 27, 31].

Despite the experimental advantages afforded by cultivation from gemmules, freshwater sponges currently pose an interpretive challenge for microbiome research. Sequencing studies report that, like their marine counterparts, freshwater sponge species harbor bacterial assemblages distinct from the surrounding environment [32–43], and these associations are often regarded as stable. For example, Paix and colleagues [33] conducted a microbiome sequencing survey across the development of gemmules in the freshwater sponge, *Spongilla lacustris*, and concluded that bacterial symbionts are transmitted across dormancy on the surface and within the interior of gemmules, in addition to being acquired from the environment during development. However, molecular evidence for a stable, resident microbiome is challenging to reconcile with legacy imaging data. Specifically, an electron microscopy study of *S. lacustris* found that the mesohyl was largely devoid of freely distributed bacteria, with most bacterial cells being observed inside phagosomes and lysosomes—consistent with ongoing phagolysosomal digestion rather than stable intercellular residence [15].

This apparent discrepancy between sequencing and imaging highlights a broader challenge imposed by sponge anatomy and feeding biology. Sponges are highly porous filter feeders with water-facing surfaces, substrate-facing surfaces, and canal/aquiferous systems that are perpetually exposed to environmental microbes (Fig. S1). The aquiferous system acts to concentrate bacterial prey, so microbes are abundant in luminal spaces and within phagocytic vacuoles at all stages of intracellular trafficking (i.e., feeding). As a result, bulk-sequencing of excised sponge tissue captures a mixture of surface-associated, luminal, and intracellular bacteria, in addition to any stable symbionts in the mesohyl. This obscures the population that is typically implied by the term “sponge microbiome” and necessitates spatially resolved imaging.

Here, we combine confocal microscopy and fluorescence in situ hybridization (FISH) to address where sponge-associated bacteria actually reside, how they are transmitted across dormancy in the freshwater sponge *Ephydatia muelleri*, and what transmission patterns account for the apparent stability of the microbiome. We find no evidence for bacterial persistence within the interior of gemmules, and little evidence for a dense bacterial community inhabiting the mesohyl. Instead, we find that bacteria are largely excluded from interior spaces by a host-secreted external matrix, lined by a continuous epithelium that forms a physical barrier to the substrate and envelops spicules and gemmules. This external, host-derived matrix supports dense microbial communities in both lab cultures and field-collected specimens, suggesting the bacterial signal detected by bulk-sequencing in freshwater sponges likely arises, at least in part, from this previously overlooked niche, and not solely from symbiotic communities residing within the mesohyl.

## Materials and Methods

### Gemmule Collection

We collected *Ephydatia muelleri* gemmules in October 2021 from “upper” Red Rock Lake, Boulder County, Colorado, USA (40.0802, −105.543), a small unofficially named body of water several hundred meters southwest of Red Rock Lake. “Sponge wool” (residual matrix) and embedded gemmules were removed from rocks, placed into 1-L screw-cap containers on ice, and transported to the laboratory where they were stored long-term in the dark at 4 °C.

### Surface-sterilization of *E. muelleri* gemmules

For surface-sterilization of *E. muelleri* gemmules, all procedures were performed under aseptic conditions in a laminar-flow hood, which was pre-sterilized with 70% ethanol and ≥30 minutes of UV irradiation prior to treatment of gemmules. 500 mL of collected lake water was filtered through 0.22 µm PES, 90 mm filter (CellTreat, Cat. no.: 50-202-045) via a vacuum prior to being added to the laminar flow hood. Sterilized lake water was then additionally filtered by a syringe-driven 0.22 µm PES, 30 mm filter (CellTreat, Part no.: 229747) into 50 mL aliquots. 4 mL of sterilized lake water was placed into 35 mm uncoated coverslip-bottom dishes (MatTek, Part no.: P35G-1.5-10-C) and these dishes received an additional 30 minutes of UV irradiation prior to being plated with 3x surface-sterilized gemmules, which aggregate to form a single sponge individual (detailed below).

Prior to surface-sterilization, >100 gemmules were removed from cold storage, selected for integrity using a dissection microscope, and transferred to a 40 µm cell strainer placed in a 6 cm Petri dish containing ∼15 mL sterilized lake water. Following the protocol of Rozenfeld and Curtis [25], gemmules were surface-sterilized in a sterile laminar-flow hood by placing the cell strainer containing gemmules directly in 1% final concentration hydrogen peroxide for 5 minutes, immediately followed by a 2-minute treatment in 2% final concentration sodium hypochlorite (NaClO), diluted from household bleach (8.25% NaClO). The cell strainer containing surface-sterilized gemmules was then rinsed thoroughly and 3x gemmules were plated in each coverslip-bottom dish containing 4 mL sterilized lake water. Of note, these gemmules are clonal, deriving from the same naturally occurring adult, and aggregate to form a single sponge individual upon hatching. To evaluate the presence of microbes on the surface and/or interior of the gemmules, sponges were reared from gemmules subjected to three conditions. The first condition was sponges reared from untreated gemmules in untreated lake water, representing natural conditions. The second condition was sponges reared from untreated gemmules in sterilized lake water, to select for gemmule-associated microbial communities. The third condition was sponges reared from surface-sterilized gemmules in sterilized lake water, to select exclusively for microbial communities associated within the gemmule interior. For all three conditions, gemmules were transferred to coverslip-bottom dishes and allowed to develop in respective media for 6–8 days in the dark at room temperature (25 °C). 6x replicates from each condition (3x gemmules/1x sponge individual per well, 18x gemmules/6x sponge individuals total) were fixed, stained, and imaged (detailed below) to evaluate for the presence of microbial/bacterial communities. No antibiotics were used at any stage in this work.

### Aquaria and husbandry for long-term growth

We developed aquaria to evaluate for evidence of long-term and transient “horizontal acquisition” of associated bacteria in *E. muelleri*. This system provided the conditions which allowed for the maintenance of all long-term (≥100 days) sponge specimens used in confocal imaging and FISH analyses.

A freshwater planted aquarium was established using commercial substrate (Fluval shrimp stratum) and planted with commercially available freshwater aquatic plants. To approximate natural environmental conditions, the aquarium was regularly seeded with water, sediment, and detritus collected from upper Red Rock Lake, the same source location for gemmules and tissue samples. After equilibration for several weeks, red cherry shrimp (*Neocaridina* spp.) were introduced to promote nutrient cycling, and many other adult forms of macroinvertebrates (e.g., *Hydra* spp., snails, and leeches) from the lake spontaneously established and proliferated in the tank during this time.

The aquarium was intermittently supplemented with a range of nutritional sources, including commercial invertebrate and algal feeds, phytoplankton suspension, fish flakes, and yeast extract. A full-spectrum LED aquarium light (Hygger NatureLite) was set to a 24-hour automatic lighting cycle. Water flow was provided by a multi-stage clip-on power filter (Fluval AC50); water changes were performed approximately once per month using reverse-osmosis water. Trace ions corresponding to 1x M-Medium [44] (1 mM CaCl₂·H₂O, 0.5 mM MgSO₄·7H₂O, 0.5 mM NaHCO₃, 0.05 mM KCl, 0.25 mM Na₂SiO₃·9H₂O) were added occasionally as needed. No chemical filtration or antimicrobial treatments were applied.

To test whether the tank could support sponges long-term in general, bulk pieces of “sponge wool” (field-collected matrix embedded with >50 gemmules) were placed on various substrates in the tank (rocks, branches, etc.). After achieving successful maturation and establishment of sponges in the aquarium (see Supplementary Videos S1 & S2), we then reared mature sponges on coverslip-bottom dishes. Here, 4 mL of untreated aquarium water was placed in coverslip-bottom dishes and 3x untreated gemmules per dish were allowed 6–8 days to develop and attach to the coverslip, again aggregating to form a single sponge individual. Myriad sponge individuals, established on coverslip-bottom dishes and all deriving from the same clonal gemmule stock, were then transferred to fish breeding/isolation boxes placed directly in the aquarium, where they were left to mature long-term. For FISH and generic microscopy, mature gemmule-hatched sponges were removed and fixed and stained on the bench top (detailed below).

### Microscopy and sample preparation

Bright field images of sponges were obtained on either an inverted scope (Olympus IX51) or a dissecting scope (Olympus SZ61). For general tissue staining of samples for confocal microscopy, fixation was performed with 4% formaldehyde in 1 mL 50% ethanol for 30 minutes, adding 1 mL of 100% ethanol every 15 minutes up to 1 hour total, as initial treatment with high ethanol concentrations compromised the integrity of the epithelium lining the external matrix. Fixation was followed by incubation with 1% bovine serum albumin (BSA) in 1x PBS + 0.1% Tween 20, followed by staining with Hoechst 33342 (100 µg mL⁻¹) and phalloidin-Alexa Fluor 568 (final concentration 3 U mL⁻¹). Samples were imaged on an Olympus IX83 confocal microscope using standard optical sectioning.

### Fluorescence in situ hybridization (FISH)

FISH largely followed a previously described protocol [45]. A near-universal bacterial probe (EUB338; 5′-GCT GCC TCC CGT AGG AGT-3′) was synthesized and conjugated to Alexa Fluor 488 (Integrated DNA Technologies), and resuspended to 100 µM in molecular-biology-grade water. Prior to fixation, live samples were incubated for 15 minutes in CellBrite 555 and washed with sterile lake water.

The gradual locomotion of both marine and freshwater sponges has been previously reported [46], which drastically reduced the amount of viable mature sponges for FISH microscopy, as most aquarium transplants of *E. muelleri* “crawled” off of the glass coverslip during maturation in the aquarium. Three aquarium-reared, ∼3-month-old sponge individuals were selected and examined for bacterial localization with FISH using the EUB338 eubacterial probe: two “non-starved” individuals and one “starved” individual. The “starved” individual was transferred to sterilized lake water for 4 days prior to fixation and sample preparation.

Samples were fixed in 4% formaldehyde with phalloidin-Alexa Fluor 568 (3 U mL⁻¹) for 1 hour, then replaced with 4% formaldehyde and incubated for an additional 3 hours. After washing, samples were incubated in 2x SSC containing 10% pre-hybridization buffer for 2 hours, then hybridized overnight (∼16 hours) at 37 °C with EUB338 probe diluted 1:100 in hybridization buffer. Post-hybridization washes consisted of a 10-minute incubation in pre-hybridization buffer, one 15-minute wash in 2x SSC, and two 15-minute washes in 0.2x SSC. Samples were counterstained with Hoechst (100 µg mL⁻¹) for 15 minutes, washed, mounted, and stored at 4 °C prior to imaging.

Negative controls included samples processed without the probe, which showed no detectable fluorescence signal. Multiple confocal fields (≥5) were acquired from all three sponge individuals; and representative images were selected to illustrate consistent spatial patterns observed across imaging fields for each individual. Importantly, the EUB338 probe was designed to target a highly conserved region of the bacterial 16S rRNA gene (present in ∼97% of bacterial 16S rRNA sequences), but has been shown to be insufficient for detection of *Planctomycetales* and *Verrucomicrobia*, and some green non-sulfur bacteria [47].

### Field-collected specimens

We collected ten small (∼0.2 mm^2^) tissue sections from two *Ephydatia muelleri* individuals (5 tissue sections per individual, 10 total) in July 2025 from “upper” Red Rock Lake, Boulder County, Colorado, USA (40.0802, −105.543), the same lake where all gemmules in this study were also collected. Tissue sections were fixed directly in the field in 1.5 mL tubes containing 500 µL 50% EtOH + 4% formaldehyde for 30 minutes, adding 500 µL of 100% EtOH every 15 minutes up to an hour. The fixative was then removed and sections were washed in 3% BSA, and subsequently allowed to rest in 1.5 mL 3% BSA for ∼2 hrs while being transported back to the University of Denver. Remaining sample preparation and staining was identical to the general tissue staining methods (detailed above).

## Results

### Evidence against seasonal bacterial carryover within *E. muelleri* gemmules

Two recent studies applied bulk microbiome sequencing techniques to evaluate the diversity of bacterial communities within the freshwater species, *E. muelleri* and *S. lacustris* [32, 33]. Both groups reported bacterial assemblages that were distinct from the surrounding environment. Paix and colleagues [33] further examined bacterial communities associated with untreated and surface-sterilized gemmules, as well as early developmental stages of *S. lacustris*, reporting reproducible and specific bacterial assemblages associated with the gemmule coat matrix, as well as within the gemmule interior, which they refer to as “vertical transmission.” However, gemmules are dormant, asexual propagules rather than sexual offspring, so vertical transmission in this context describes persistence of bacteria across seasonal dormancy.

To directly test whether *E. muelleri* retains bacterial communities within the interior of gemmules across periods of dormancy, we surface-sterilized gemmules and reared them in sterile medium (sterilized lake water) without the addition of antibiotics. In line with previous reports [25], these treatments had no apparent effect on sponge development (Fig. 1A). Surface-sterilization of gemmules reduced the hatch rate by approximately 30%; 9 of 30 treated gemmules failed to develop whereas all untreated gemmules hatched successfully (36/36).

**Fig. 1.**
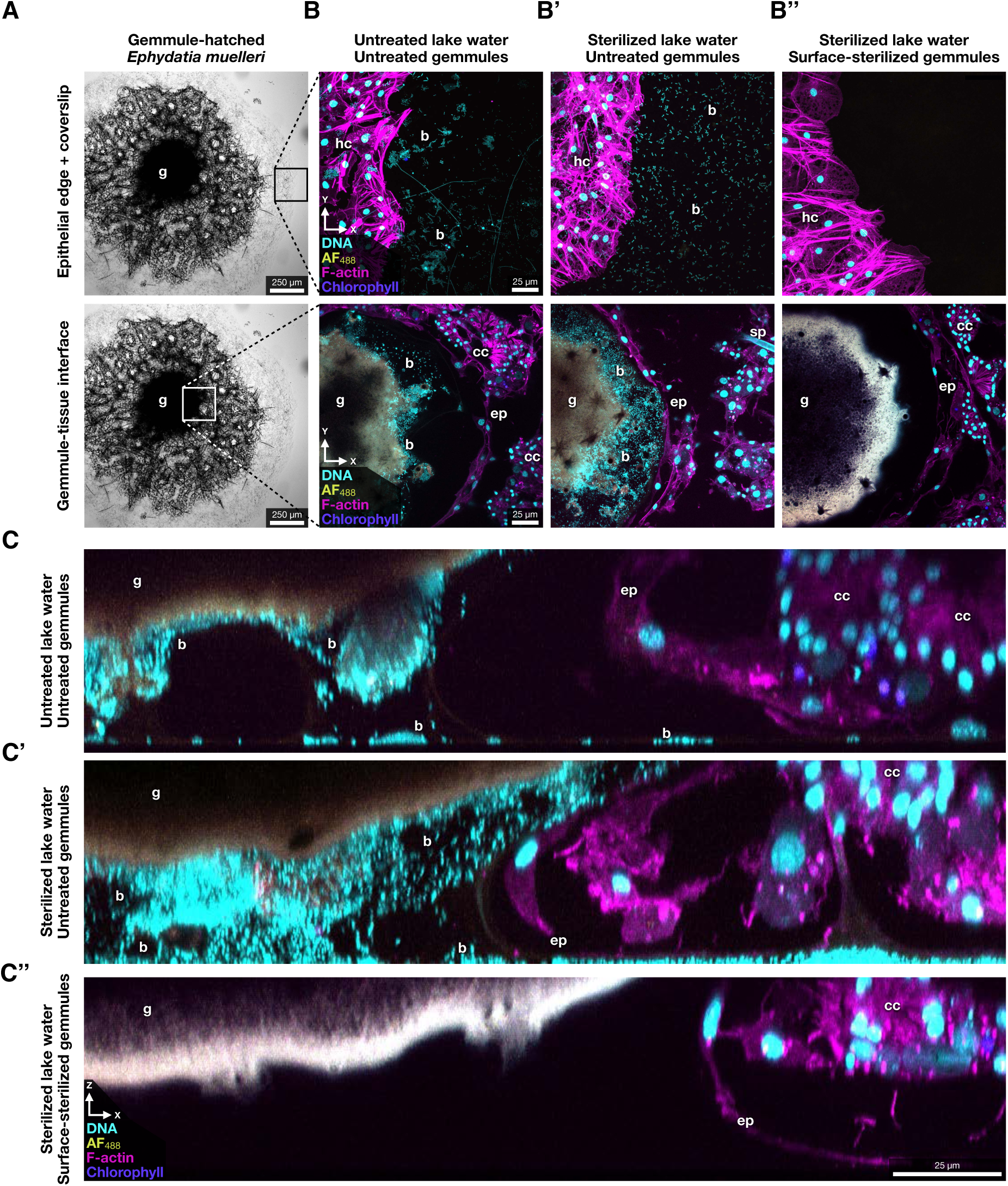
Stable bacterial communities are not present within gemmules. **(A)** Representative light micrograph of an *E. muelleri* juvenile hatched from a single gemmule (scale bar = 250 µm). **(B–B″)** Representative images of the epithelial edge and coverslip (*top*) and gemmule-tissue interface (*bottom*) of sponge individuals reared from three clonal gemmules under different microbial exposure conditions. **(B)** Sponges derived from untreated gemmules in natural lake water show diverse bacteria associated with the coverslip and at the gemmule-tissue interface. **(B′)** Sponges derived from untreated gemmules in sterilized lake water show more morphologically uniform bacteria on the coverslip and at the gemmule-tissue interface. **(B″)** Sponges derived from surface-sterilized gemmules in sterilized lake water show no observable bacteria on the coverslip or at the gemmule-tissue interface. **(C–C”)** Respective orthogonal projections of the gemmule-tissue interface shown in the bottom panels of (B–B”) highlight epithelium separating host tissues from the gemmule. (b:bacteria; cc: choanocyte chamber; ep: epithelium; g: gemmule; hc: host cells; sp: spicule; cyan = DNA; magenta = F-actin; scale bar = 25 µm)

Three conditions were compared using confocal microscopy: untreated gemmules in natural lake water, untreated gemmules in sterile medium, and surface-sterilized gemmules in sterile medium (Fig. 1). For each condition, six sponges, each grown from three clonal gemmules, were evaluated. As expected, all sponges (6/6) grown from untreated gemmules in natural lake water exhibited abundant, morphologically diverse bacteria, observed as a nascent biofilm on the coverslip, as well as at the gemmule-tissue interface (Fig. 1B–C). In comparison, all sponges (6/6) grown from untreated gemmules in sterilized lake water also exhibited microbial communities on the coverslip and at the gemmule-tissue interface, though these bacterial populations were more morphologically uniform (Fig. 1B’–C’). In contrast, no observable bacteria were detected in five of six (5/6) sponges grown from surface-sterilized gemmules in sterilized lake water, on the coverslip or at the gemmule-tissue interface (Fig. 1B”–C”). In the single sponge exhibiting incomplete surface-sterilization of gemmules, only a few isolated bacterial cells were found on the coverslip or at the gemmule surface.

These results indicate that *E. muelleri* does not maintain a persistent internal bacterial community within gemmules through periods of dormancy. Rather, bacteria associated with gemmules remain largely excluded from the mesohyl by a continuous epithelium. Bacterial signatures in other studies are thus more consistent with post-hatching environmental acquisition or persistence of bacterial communities on the external gemmule coat, rather than sustained bacterial populations being transmitted within the gemmule interior across dormancy. (Of note, intracellular *Chlorella*-like symbiotic algae are still present in cultures of surface-sterilized gemmules reared in sterilized lake water.)

### Horizontal acquisition and feeding artifacts in mature sponges

We next examined whether *E. muelleri* acquires a symbiotic bacterial community transiently from the environment (i.e., by horizontal acquisition). Gemmule-hatched sponges only remain viable when unfed in sterile medium for ∼12 days post-plating (dpp) (Fig. S2A). For longer-term growth, gemmule-hatched sponges were established on coverslip-bottom dishes and, following attachment, were maintained in commercial fish-breeding boxes in a planted aquarium designed to mimic natural lake conditions (see Methods) (Fig. S2B–C; Supp. Videos SV1 & SV2). After ∼100 days in the aquarium, three mature sponges were analyzed for the presence of bacteria by FISH using the nearly universal eubacterial 16S rDNA probe, EUB338 [48].

Confocal imaging of mature, aquarium-reared sponges revealed a previously undescribed feature of sponge anatomy: an external matrix at the basal substrate interface. This matrix forms a continuous barrier that physically separates environmental biofilms from the basal epithelium (basopinacoderm) (Fig. 2A–B, Fig. S3). The autofluorescence of this basal matrix resembled that of the gemmule coat (Fig. 1 B–C”) and imaging revealed abundant, uniformly-sized, rod-shaped bacteria (Fig. 2B’). In contrast, within the mesohyl we detected only sparse EUB338-positive puncta that were morphologically distinct from environmental and matrix-associated bacteria in both size and shape. These puncta were typically within or on the surface of cells, not freely distributed in the mesohyl (Fig. 2B’’). Although this could reflect fixation artifacts of a highly aqueous extracellular matrix, the absence of comparable clustering of migratory, mesohyl-resident host cells argues against this explanation alone.

**Fig. 2.**
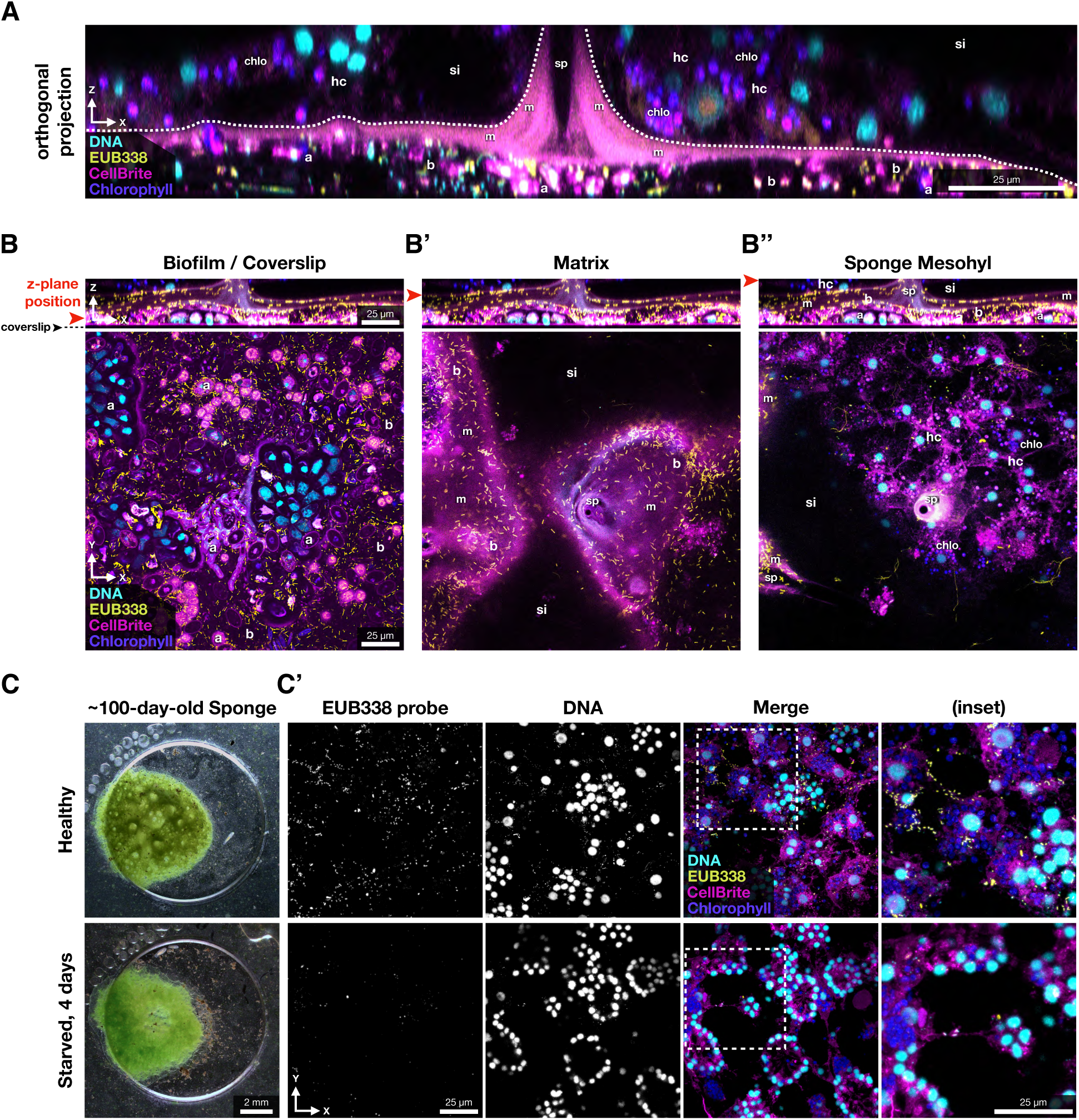
FISH with a eubacterial 16S rRNA probe and long-term rearing reveals novel physiology in *E. muelleri*. **(A)** Orthogonal projection of fluorescent in-situ hybridization of a eubacterial probe (EUB338[48]) showing an autofluorescent collagenous matrix secreted at the basal substrate attachment interface. This matrix forms a physical barrier separating the microbe-rich biofilm (below white dashed line) from the sponge interior and host cells (above dashed line). A siliceous spicule is embedded within this matrix. **(B–B”)** A single confocal scan of a ∼100-day-old, aquarium-reared *E. muelleri* individual imaged with the EUB338 probe. *Top*, orthogonal projection of the full stack; arrows indicate representative planes shown in B–B″. **(B)** A stack near the coverslip shows a dense biofilm beneath the sponge, covered by the same matrix depicted in A. **(B′)** A stack highlighting the basal collagenous matrix, which is occasionally colonized by uniformly sized, rod-shaped bacteria. **(B″)** A stack highlighting the sponge interior and host cells, where EUB338-positive puncta are markedly smaller than bacteria observed within the biofilm or basal matrix. **(C–C’)** Starvation experiments suggest that most EUB338-positive puncta within the sponge interior arise from the canonical feeding pathway. **(C)** Representative images of a healthy ∼100-day-old sponge (*top*) and a sponge starved for 4 days by removal from the aquarium (*bottom*). **(C′)** Confocal stacks of the choanoderm of a healthy sponge (*top*) and a sponge starved for 4 days (*bottom*) using FISH with the EUB338 probe highlighting a sharp reduction in EUB338-positive puncta after 4 days of starvation. (a: algae; b: bacteria; hc: host cells; m: collagenous matrix; si: sponge interior; sp: spicule; DNA = cyan; EUB338 probe = yellow; F-actin / 568 nm autofluorescence = magenta; Chlorophyll autofluorescence at 640 nm = blue; scale bar = 25 µm)

We therefore sought to further address the nature of these internal EUB338-positive puncta, as the biological origin of these signals was uncertain and could reflect intercellular bacteria, partially degraded bacterial material derived from feeding, or other transient microbial remnants. To assess whether their abundance was linked to nutritional status, we “starved” a single ∼100-day-old sponge by transferring it to filter-sterilized lake water for 4 days (Fig. 2C). Following starvation, the internal EUB338-positive puncta were largely eliminated (Fig. 2C’). Previous studies have suggested that sponges may metabolize symbiotic bacteria under nutrient limitation [49], raising the possibility that internal bacterial material was metabolized during this period of starvation. The small size, sparse distribution, and rapid loss of these puncta upon starvation are inconsistent with the presence of a dense, stable, or spatially extensive bacterial community within the mesohyl of *E. muelleri*. Critically, however, hybridization with the EUB338 probe denotes these puncta as plausible substrates for 16S rRNA amplicon sequencing, irrespective of their functional significance to the sponge host.

### Bacteria colonize the external matrix that embeds siliceous spicules

In addition to forming a continuous barrier at the substrate-biofilm interface, the external matrix was also found to surround and embed siliceous spicules, consistent with previous descriptions [29] of the skeletal lattice of freshwater sponges (Fig. 2A–B”). Like the matrix coating gemmules and lining the substrate, this spicule-associated sheath contained discrete pockets of bacteria in aquarium-reared sponges. Notably, these bacterial populations were again separated from the mesohyl by a continuous epithelium (Fig. 3A).

**Fig. 3.**
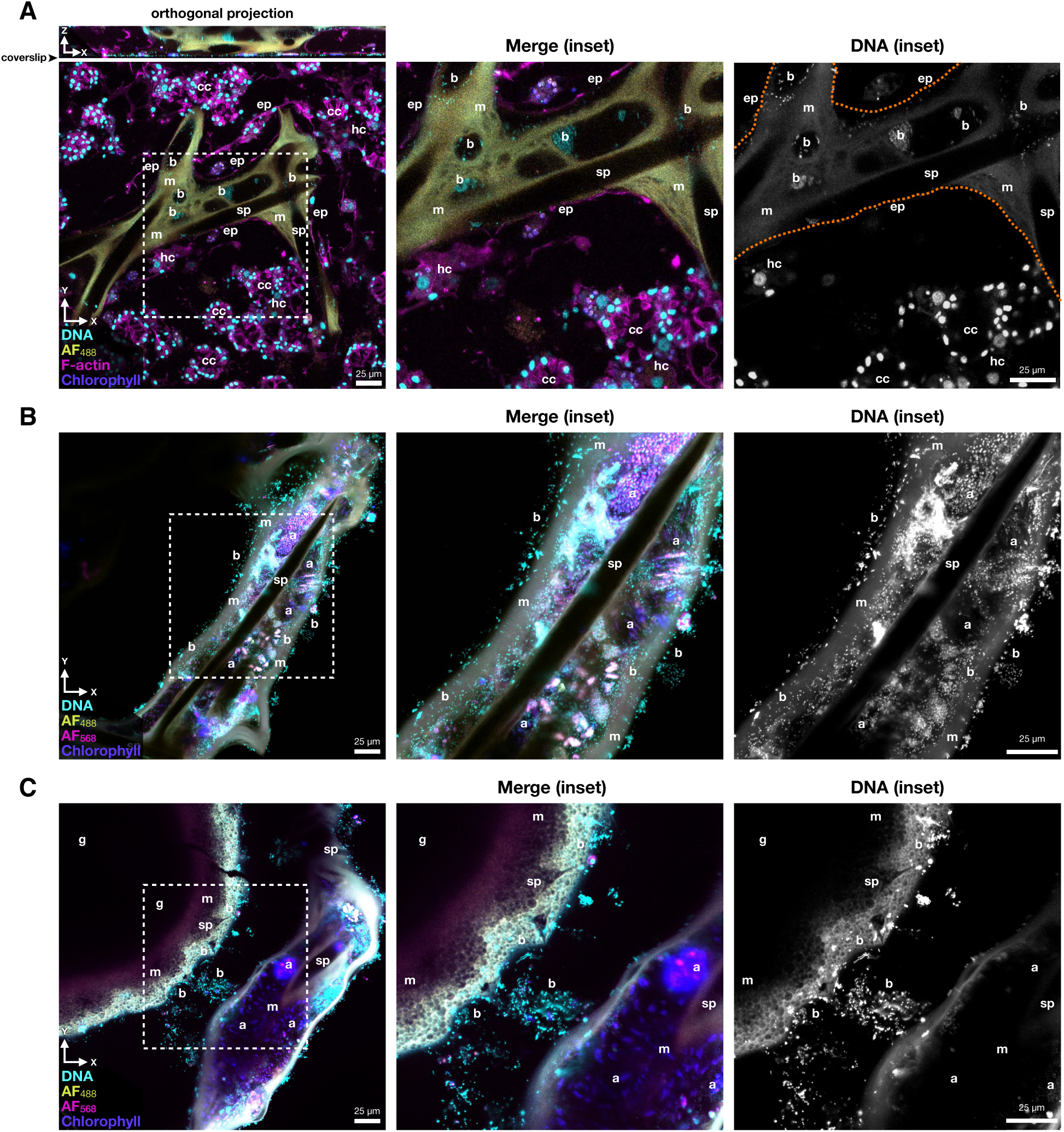
Confocal imaging of *E. muelleri* collagenous skeletal matrix reveals its colonization by microbes. **(A)** Confocal stack of a gemmule-hatched E. muelleri showing a largely bacteria-free mesohyl and a collagenous skeletal matrix that retains bacterial colonies. Orange dashed lines delineate the epithelial boundary separating the collagenous skeletal matrix from the sponge body cavity. **(B)** Confocal micrograph of field-collected skeletal matrix from naturally occurring *E. muelleri* showing a diverse microbial community, including bacteria and photosynthetic microeukaryotes, residing within the matrix. **(C)** Confocal micrograph of field-collected skeletal matrix containing an unhatched gemmule embedded within the matrix, highlighting bacterial colonization of the collagenous gemmule coat and a dense microbial community within the adjacent skeletal matrix. (a: algae; b: bacteria; cc: choanocyte chambers; ep: epithelium; g: gemmule; m: matrix; sp: spicule; DNA = cyan; 488 nm autofluorescence = yellow; F-actin = magenta in (A); 568 nm autofluorescence = magenta in (B–C); Chlorophyll autofluorescence at 640 nm = blue; scale bar = 25 µm)

To determine whether these observations from aquarium-reared sponges also reflect conditions in nature, we examined the spicule-associated external matrix in field-collected specimens undergoing tissue regression and dormancy. Bare skeletal matrix, with no overlying sponge tissue, was densely colonized by diverse microbial communities, including bacteria and photosynthetic microeukaryotes (Fig. 3B). Similarly, bacteria were abundant on the surfaces of field-collected gemmules (Fig. 3C). Together, these observations suggest that the spicule-associated matrix functions less like an internal skeleton and more as a substrate over which regenerating sponge tissue propagates [50]. During dormancy, both the gemmule coat and spicule-associated matrix are exposed to the environment, and can be colonized by microbes. When conditions improve, tissue emerging from gemmules spreads over this pre-existing, microbe-colonized skeletal scaffold, while a defined epithelium maintains separation between the matrix and the mesohyl.

We tested this hypothesis in two ways. First, we followed a single *E. muelleri* individual for 549 days in the aquarium. During this period, the sponge underwent multiple rounds of gemmulation and regeneration, repeatedly building new tissue over the residual spicule-associated matrix (Fig. S4A). Imaging at day 549 revealed a consistent spatial pattern: dense microbial communities were present at the basal substrate interface (Fig. S4B) and on mature spicule-associated external matrix (Fig. S4C). However, newly deposited matrix around developing spicules was devoid of observable bacteria (Fig. S3D). Additional imaging of gemmule-hatched sponges further supported that newly deposited matrix does not retain any observable bacteria, irrespective of development on an established biofilm at the basal substrate interface (Fig. S5). Thus, it seems microbial colonization of the matrix occurs after matrix synthesis and deposition.

Second, we asked whether the spatial organization observed in aquarium-raised sponges also occurs in natural populations. We fixed tissue sections (n = 10 from 2 individuals) immediately in the field for evaluation by confocal microscopy. The pattern was clear: matrix and associated microbes remained separated from the host mesohyl by an epithelium (Fig. 4). The matrix surrounding large, mature spicules contained complex microbial communities (Fig. 4, inset 1); and regions containing newly deposited matrix around smaller, developing spicules lacked detectable microbes (Fig. 4, inset 2). This was consistent with patterns seen in aquarium-reared sponges. We cannot distinguish whether the myriad Hoechst-positive puncta associated with sponge cells represent digestive vacuoles, bacterial prey, or mesohyl-resident symbionts—though they appear to be primarily intracellular.

**Fig. 4.**
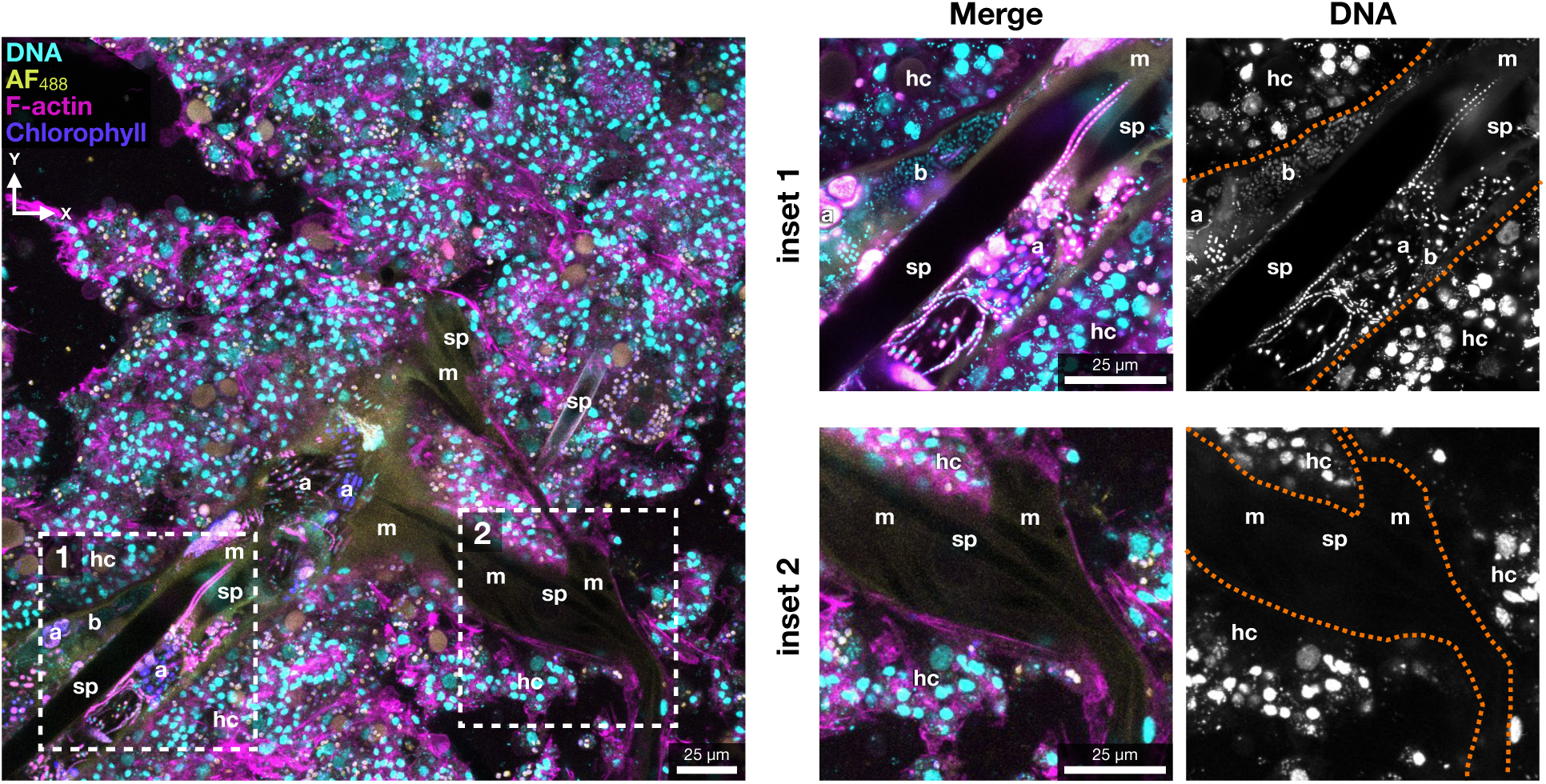
Microbial communities reside within the skeletal matrix of tissue sections from field-collected *E. muelleri*. Confocal stack of a tissue section from naturally occurring *E. muelleri* shows a robust microbial community, consisting of both bacteria and photosynthetic microeukaryotes, residing within the portion of matrix surrounding large spicules **(inset 1)** and a relatively microbe-free portion of matrix surrounding smaller, newly-developing spicules **(inset 2)**. In both cases, sponge host cells and mesohyl remain physically separated from the matrix. Orange dashed lines delineate epithelium physically separating the collagenous skeletal matrix from the sponge body interior. (a: algae; b: bacteria; hc: host cells; m: matrix; sp: spicule; DNA = cyan; 488 nm autofluorescence = yellow; F-actin = magenta; Chlorophyll autofluorescence at 640 nm = blue; scale bar = 25 µm)

Together, these data support a model where the spicule-associated external matrix of *E. muelleri* is primarily colonized by bacteria during periods of dormancy, and subsequently overgrown by regenerating host tissue. Matrix-associated bacteria and microeukaryotes remain physically excluded from the mesohyl by an epithelium. More broadly, this spatial organization provides a conceptual bridge between sequencing studies of bulk tissue and legacy imaging of the mesohyl of freshwater sponges.

## Discussion

### Re-evaluating transmission modes and mesohyl residence of microbial communities in freshwater sponges

In critical animal-microbe symbioses, microbial partners are often transmitted vertically, directly from parent to offspring, to ensure faithful inheritance across generations [51–54]. In *E. muelleri*, two recent studies reported enrichment of *Sediminibacterium* sequences in both gemmules and adult tissues collected from Sooke River, Canada [32, 55], a pattern interpreted as possible evidence of vertical transmission [32]. Similarly, Paix and colleagues [33] reported vertical transmission of epibacteria on the external gemmule coat, as sponges hatched from surface-sterilized gemmules (following Leys et al. [56]) were found to have a less diverse microbiome compared to untreated gemmules. Sequencing of these surface-sterilized gemmules was reported as evidence of a microbial community being maintained within the gemmule interior. On the basis of these and other studies [33, 36, 57–59], the transmission and maintenance of a stable microbiome across dormancy in freshwater sponge gemmules has recently been reiterated in a broader review context [30]. As in marine species, the current consensus on bacterial microbiome transmission modes in freshwater sponges supports “mixed-mode” transmission (i.e., a combination of vertical and horizontal transmission) [32, 33, 49, 60, 61].

We directly tested the mixed-mode transmission model in *E. muelleri* using spatially resolved imaging across dormancy, early development, and mature tissues. In contrast to sequencing-based studies, we found no evidence that bacterial populations are maintained within the interior of gemmules after robust surface-sterilization (Fig. 1B–C”). We next examined whether bacteria accumulate within the mesohyl over longer time scales following environmental exposure by imaging mature, aquarium-reared individuals using FISH. In these specimens, EUB338-positive puncta within the sponge mesohyl were consistently smaller than those seen in biofilms or within the matrix. These puncta were generally intracellular, or closely apposed to the surface of host cells, in contrast to the freely distributed bacteria inhabiting the mesohyl of some marine species [8–14].

The biological origin of these signals remains uncertain. One possibility is that they represent bacteria or bacterial debris associated with feeding and intracellular processing, including partially degraded, prey-derived remnants, or even residual rRNA or DNA within phagolysosomal compartments. Alternatively, they could indicate a low-abundance population of bacteria in close association with host cells in the mesohyl. To evaluate whether these puncta reflect stable association versus a feeding-contingent signal, we starved aquarium-reared sponges for 4 days in filter-sterilized lake water. Under these conditions, the internal EUB338-positive puncta were essentially eliminated (Fig. 2C’), a pattern more consistent with rapid turnover of bacterial food/debris. While it has been suggested that sponges may metabolize symbiotic bacteria during nutrient limitation [49], distinguishing selective depletion of a resident microbiome from reduced feeding-contingent signal will require approaches that directly assay bacterial viability, replication, and spatial persistence.

Several factors likely contribute to the discrepancy between our findings and prior studies. Methodologically, Paix and colleagues [33] relied on hydrogen peroxide alone for gemmule surface-sterilization, whereas earlier work reported that total removal of gemmule-surface-associated bacteria requires a combination of hydrogen peroxide and bleach treatments [25], which we applied here. In addition, amplicon sequencing is inherently more sensitive than imaging and can detect extremely rare bacteria, non-viable cells, or residual 16S rRNA/DNA that persists after sterilization. Finally, our results could reflect intrinsic biological differences between species and environmental variation; Paix and colleagues [33] examined *S. lacustris* from lowland European freshwater systems whereas we examined *E. muelleri* from high-altitude lakes in western North America. However, an independent ultrastructural study of *S. lacustris* from pre-alpine lakes in southern Germany also reported the mesohyl to be largely devoid of free bacteria, with microbial cells confined to phagosomes and lysosomes—consistent with feeding rather than intercellular residence in the mesohyl [15].

Taken together, our imaging data indicate that bacteria are not retained through dormancy within gemmules of *E. muelleri*, and that bacterial populations present within the mesohyl are relatively rare compared to those found within the external skeletal matrix. Thus, bulk-sequencing datasets in freshwater sponges are unlikely to primarily reflect stable, intercellular bacterial communities within the mesohyl. Similar imaging techniques and reasoning should be applied to assess bacterial localization and functional significance in marine sponge species.

### Recognizing the external matrix in freshwater sponges as a microbial niche bridges sequencing and spatial data

Unexpectedly, our imaging analyses revealed that *E. muelleri* produces a prominent extracellular matrix located outside the epithelial boundary that is permissive to microbial colonization. This matrix forms a continuous barrier at the basal substrate interface, in addition to embedding siliceous spicules (Fig. 2A–B”; Fig. S2). The properties of this matrix closely match classical descriptions of the collagenous “cementing substance” that anchors/coats freshwater sponge spicules and anchors gemmules [29, 62], indicating it reflects a widely deployed external matrix architecture.

Several of our observations indicate that this matrix—rather than the sponge mesohyl alone—is a credible source of bacterial signal recovered in bulk-sequencing studies. In both laboratory cultures and field-collected specimens, we found that dense and visually diverse microbial communities were consistently associated with this matrix, while a continuous epithelium maintained strict physical separation from the mesohyl (Fig. 3A; Fig. 4). In contrast, newly formed spicule matrix remained largely microbe-free (Fig. 4, inset 2; Fig. S4). Thus, it seems that opportunistic microbes colonize residual spicule-associated matrix during dormancy, when the matrix is exposed to the environment (see Graphical Abstract). Indeed, we found that this matrix harbors diverse microbial communities, even in the absence of overgrowing sponge tissue (Fig. 3B–C). Fundamentally distinct from an internal skeleton, the residual spicule matrix of freshwater sponges represents a novel microbial niche that ultimately serves as a de facto substrate for regenerative sponge tissue, and which remains physically partitioned from the host mesohyl by a continuous epithelium.

This spatial and temporal pattern suggests that microbial colonization of the matrix is secondary and environmentally driven, rather than the result of host-driven processes. However, we recognize the possibility that external matrix-associated microbes may be biologically relevant, and could confer yet-unknown effects on the sponge host. In other animals, external microbial communities often play critical protective or developmental roles. For example, in the Hawaiian bobtail squid, females inoculate the mucous coatings of their eggs with specific bacteria that inhibit biofouling and are required for canonical development [51]. In that system, microbes remain external to embryonic tissues, yet are nonetheless essential for host fitness [7]. Experimental depletion of these bacteria completely disrupts female reproductive organ development [63]. By analogy, bacteria associated with the gemmule coat or external matrix of freshwater sponges could plausibly function to prevent biofouling, produce chemical defenses, or modulate microbial succession during dormancy. Considering we observed a ∼30% reduction in hatch rate for surface-sterilized gemmules, future work should further investigate fitness contributions of gemmule-coat-associated bacteria.

Under our model, microbes associated with *E. muelleri* are spatially segregated within an external compartment rather than a host-privileged internal one. Such associations could arise through passive environmental filtering mediated by matrix chemistry—favoring microbes capable of adhering to, tolerating, or metabolizing matrix components—rather than through active host curation or coevolution. This model reframes the detection of similar bacterial taxa in sequencing datasets of freshwater sponges with the limited and inconclusive evidence for mesohyl-resident bacteria in spatial analyses. Apparent “microbiomes” in freshwater sponges—and potentially in some marine species—may therefore reflect stable occupation of host-derived extracellular niches, rather than exclusively within host tissues.

### Limitations of the study

Several limitations should be considered when interpreting these findings. First, the eubacterial 16S rRNA FISH probe used (EUB338) does not recognize all bacterial taxa, raising the possibility that some bacteria were not detected [47], though Hoechst was always used as a counterstain. Second, imaging resolution within the densely packed interior of viable gemmules limits definitive exclusion of extremely rare internal bacteria; accordingly, we relied primarily on the absence of bacteria on the coverslip and at the gemmule-tissue interface as a measure of microbial carryover through dormancy. Third, although our aquarium conditions recapitulate many features of natural freshwater habitats, they cannot fully capture the ecological complexity or seasonal variability experienced by sponges in situ, a limitation we sought to mitigate through parallel analysis of field-collected specimens. Finally, while this study establishes a novel spatial distribution of microbes, it does not directly test whether matrix-associated bacteria influence sponge physiology, fitness, or resistance to fouling. Determining the biochemical composition of the external matrix and characterizing its associated microbiota will be essential next steps.

### Conclusions

Together, our findings show that apparently stable freshwater sponge “microbiomes” do not necessarily denote internal symbioses, and that spatial context is essential for interpreting host–microbe associations. Despite a feeding mode that necessitates continuous bacterial uptake, *E. muelleri* seems to largely conform to a broader metazoan organizational principle: host tissues remain privileged by physical exclusion of environmental bacteria, while microbial interactions occur significantly in association with intracellular membrane-bound vesicles or a secreted extracellular matrix located outside an epithelial boundary. These results reframe sequencing-based surveys with ultrastructural imaging data and suggest that, at least for some freshwater sponges, microbial abundance reflects external niche construction rather than exclusively internal symbioses. Although more technically challenging, similar imaging studies should be conducted to contextualize sequence-based microbiomes in marine species as well.

## Acknowledgements

This work was supported by NSF IOS-2426160 to S.A.N.

## Supplemental Figures

**Fig. S1.**
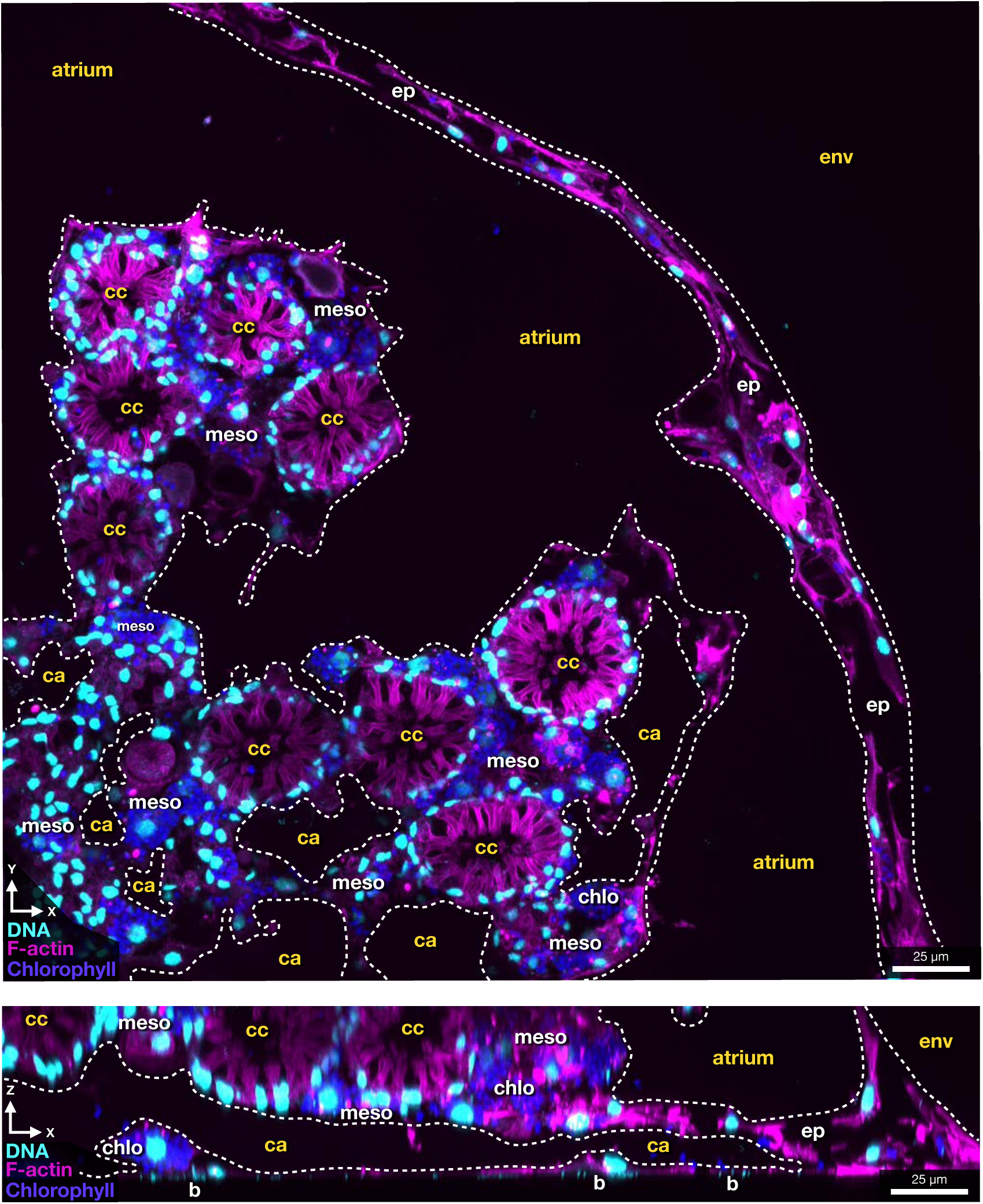
The tent epithelium of *E. muelleri* is spatially distinct from the choanoderm/mesohyl. (*Top*) Single confocal stack of an *E. muelleri* individual highlighting the tent epithelium and choanoderm/mesohyl. Environment-facing water canals are found interspersed between the two tissues. (*Bottom*) Aligned orthogonal projection of the same confocal stack highlighting the same regions. Canals also run underneath along the basal portion of the sponge choanoderm. Bacterial signatures via DNA stain are found primarily within biofilms at the coverslip, and remain largely excluded from host tissues. Labels shown in white depict internal host tissues. Labels shown in yellow depict external, environment-facing regions. (b: bacteria; ca: canal; cc: choanocyte chamber; chlo: *Chlorella*-like symbiotic algae; env: external environment; ep: epithelium; meso: mesohyl; atrium: water-filled atrium; DNA = cyan; F-actin = magenta; Chlorophyll autofluorescence at 640 nm = blue; scale bar = 25 µm)

**Fig. S2.**
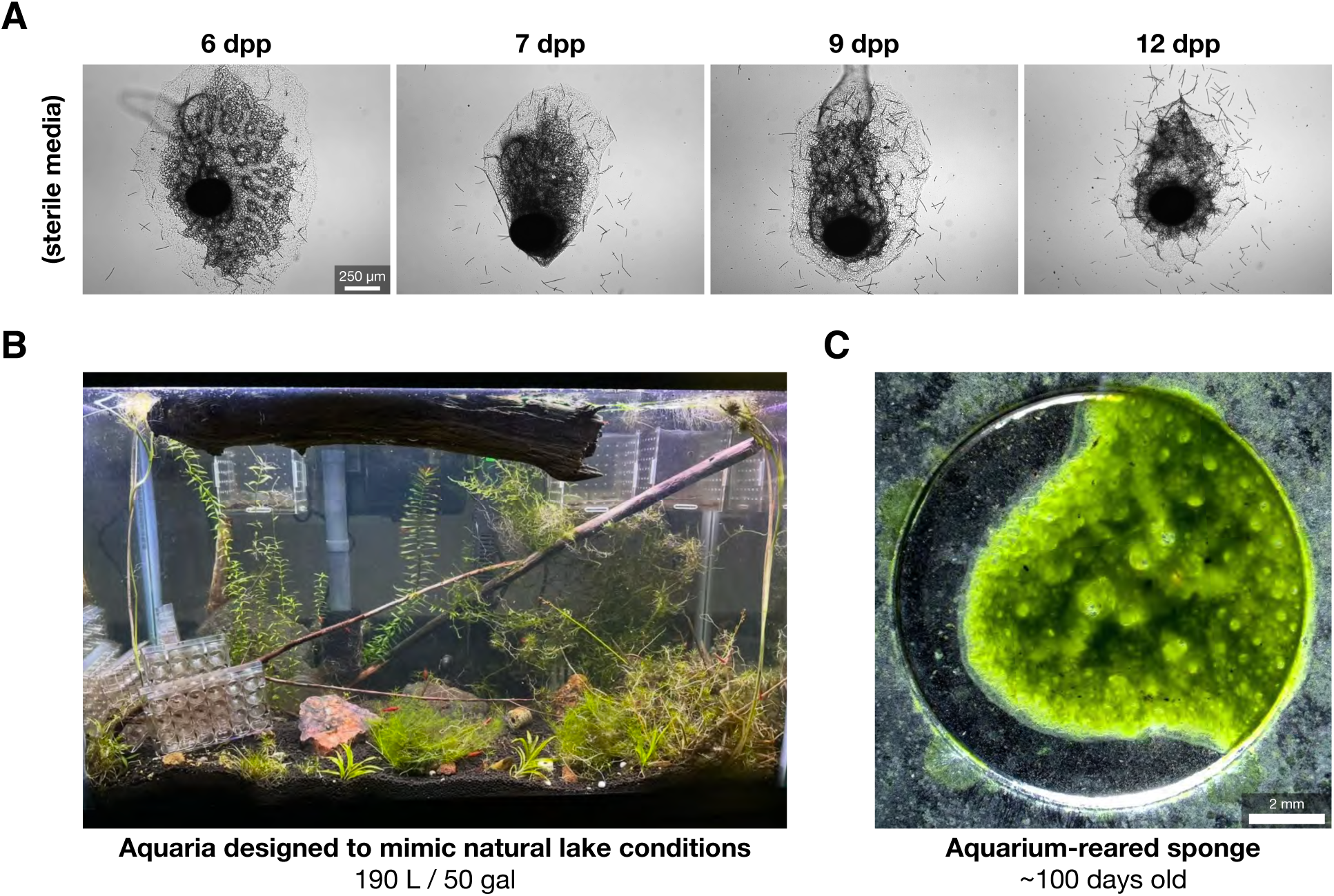
Aquaria mimicking the lake environment allows for long-term rearing of *E. muelleri*. **(A)** Representative time-course images showing viability of gemmule-hatched *E. muelleri* declines after ∼9 days post-plating (dpp) (scale bar = 250 µm) **(B)** Representative image of a ∼190 L aquarium (50-gallon) seeded with plants, soil, detritus, and water from the lake to mimic the lake environment, permitting the long-term rearing of gemmule-hatched *E. muelleri* in the laboratory. **(C)** ∼100-day-old gemmule-hatched sponge reared long-term in the aquarium on a coverslip bottom dish (scale bar = 2 mm).

**Fig. S3.**
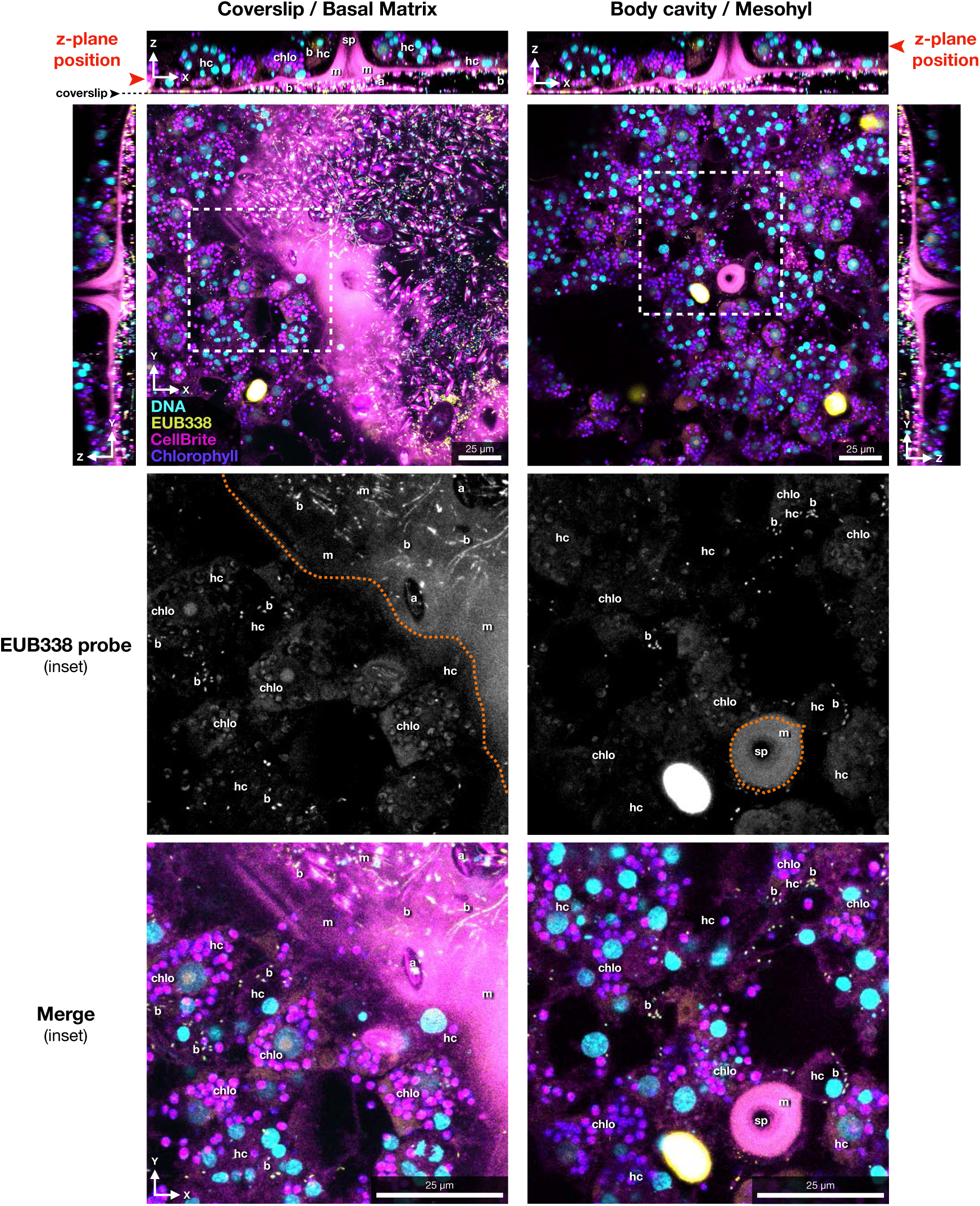
FISH with a eubacterial 16S rRNA probe and long-term rearing reveals novel sponge physiology in *E. muelleri*. Single confocal stack of a ∼100-day-old, aquarium-reared *E. muelleri* imaged with FISH using a eubacterial 16S rRNA probe (EUB338). Respective z-plane positions are shown along the top and sides and highlight the secreted basal collagenous matrix with a spicule embedded within the matrix. Stack at the coverslip (*left*) shows a defined barrier between the biofilm and host cells. Stack within the sponge body interior (*right*) shows tiny EUB338 puncta primarily being intracellular and at the cell surface of sponge host cells. (a: algae; b: bacteria; bc: body cavity; chlo: intracellular, *Chlorella*-like symbiotic algae; hc: host cells; m: collagenous matrix; sp: spicule; DNA = cyan; EUB338 probe = yellow; CellBrite / 568 nm autofluorescence = magenta; Chlorophyll autofluorescence = blue; scale bar = 25 µm)

**Fig. S4.**
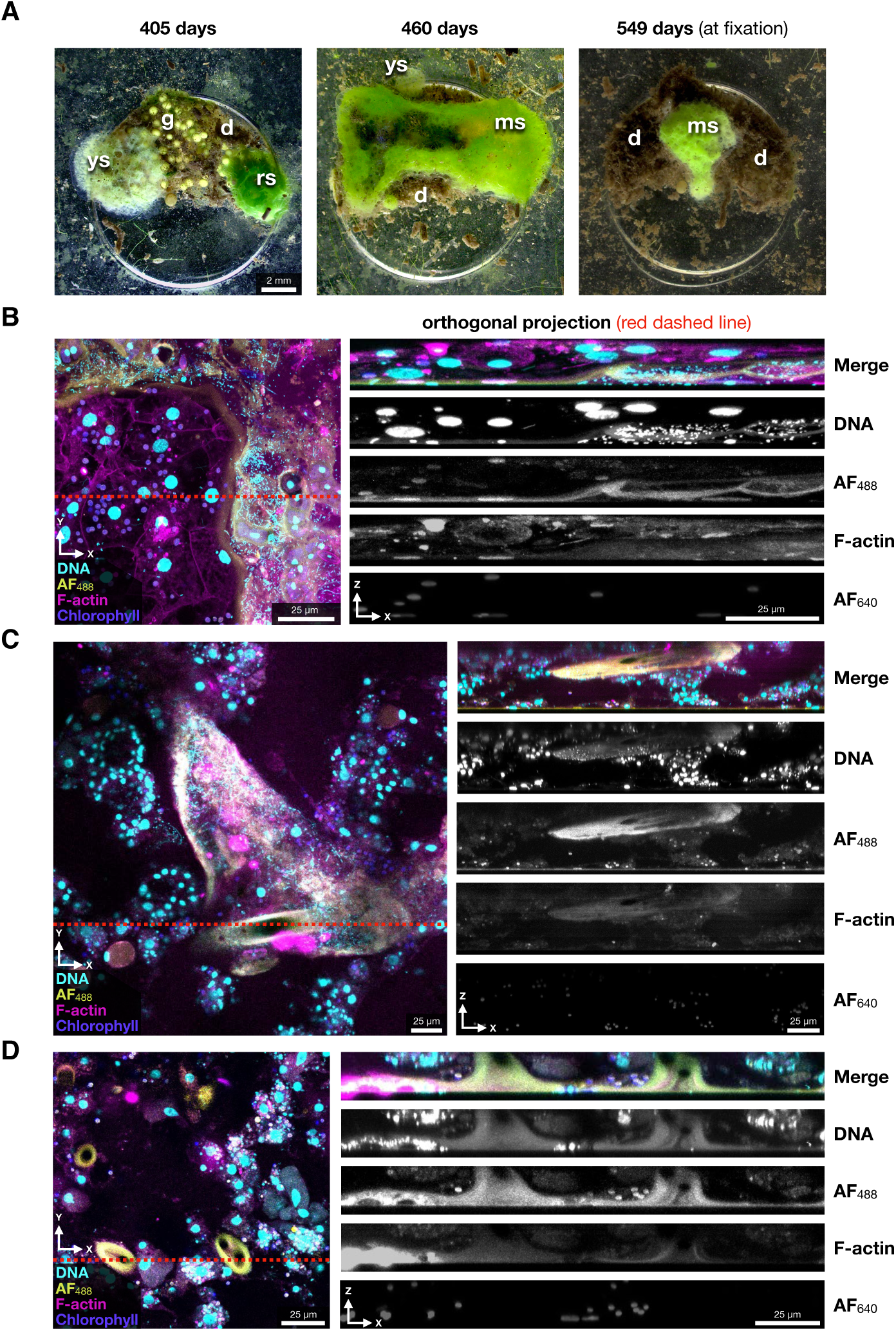
Imaging of a single, aquarium-reared sponge at day 549 supports secondary colonization of the matrix during periods of dormancy. **(A)** Representative images of a 549-day-old specimen showing various stages of asexual reproductive cycles occurring within the aquarium (scale bar = 2 mm). **(B)** Confocal stack of matrix at the basal substrate interface shows substantial microbial colonization of the matrix with the above host cells and mesohyl being relatively microbe-free. **(C)** Confocal stack of skeletal matrix of the sponge shows significant microbial colonization compared to the sponge cells and mesohyl seen around the matrix. **(D)** Confocal stack of newly developing matrix/spicules was found to retain no detectable microbial populations. (d: dead tissue; g: gemmules; ms: mature sponge; rs: regressing sponge tissue; ys: young sponge; DNA = cyan; 488 nm autofluorescence = yellow; F-actin = magenta; Chlorophyll autofluorescence = blue; scale bar = 25 µm)

**Fig. S5.**
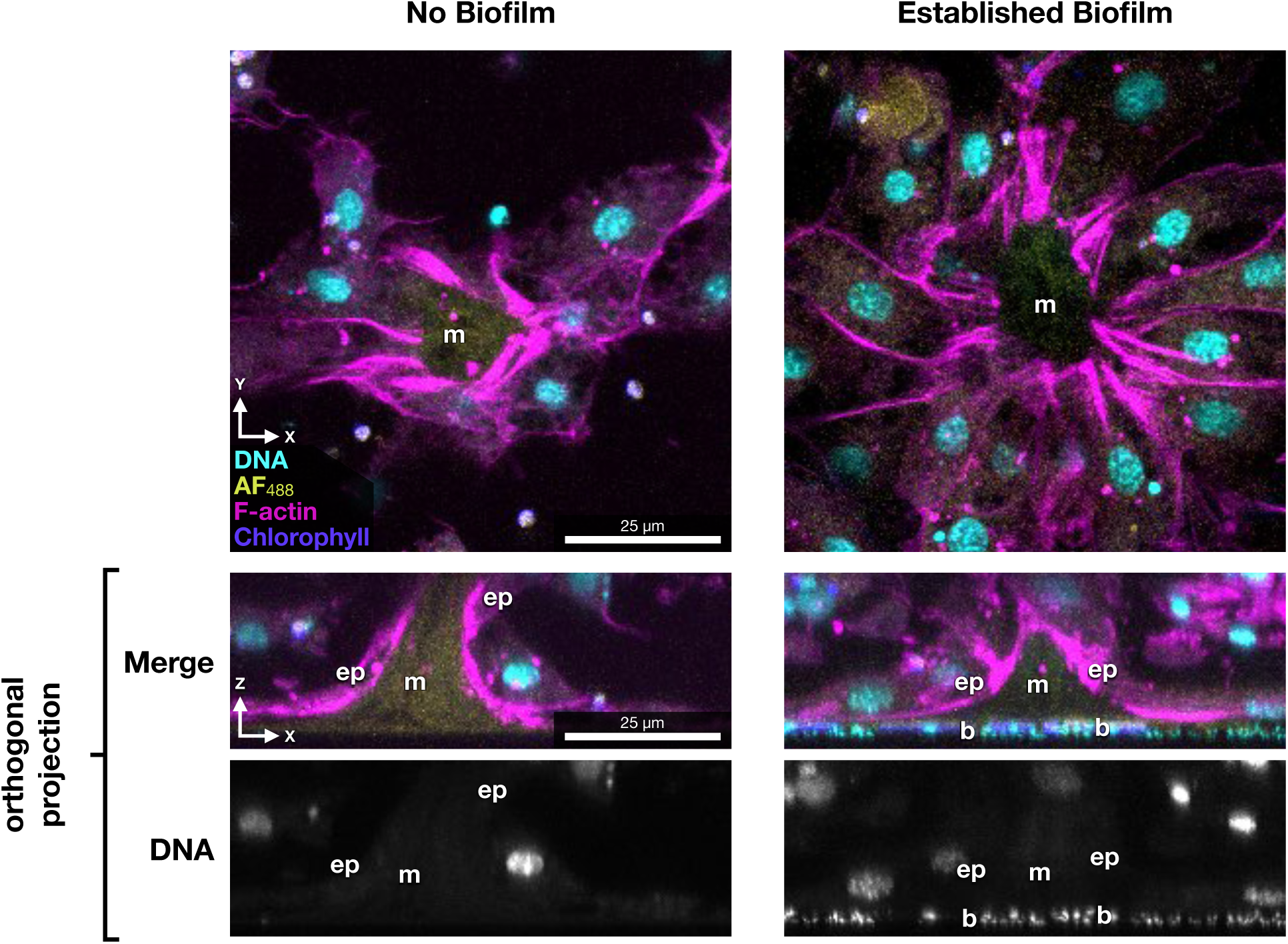
Newly developing matrix retains no detectable microbial communities, irrespective of an established biofilm. Confocal stacks of newly developing matrix/spicules were found to retain no detectable microbial populations in sponges hatched on a sterile coverslip (*left*) as well as for sponges hatched on a mature, established biofilm (*right*). (DNA = cyan; 488 nm autofluorescence = yellow; F-actin = magenta; Chlorophyll autofluorescence = blue; scale bar = 25 µm)

**Supplementary Video 1:**
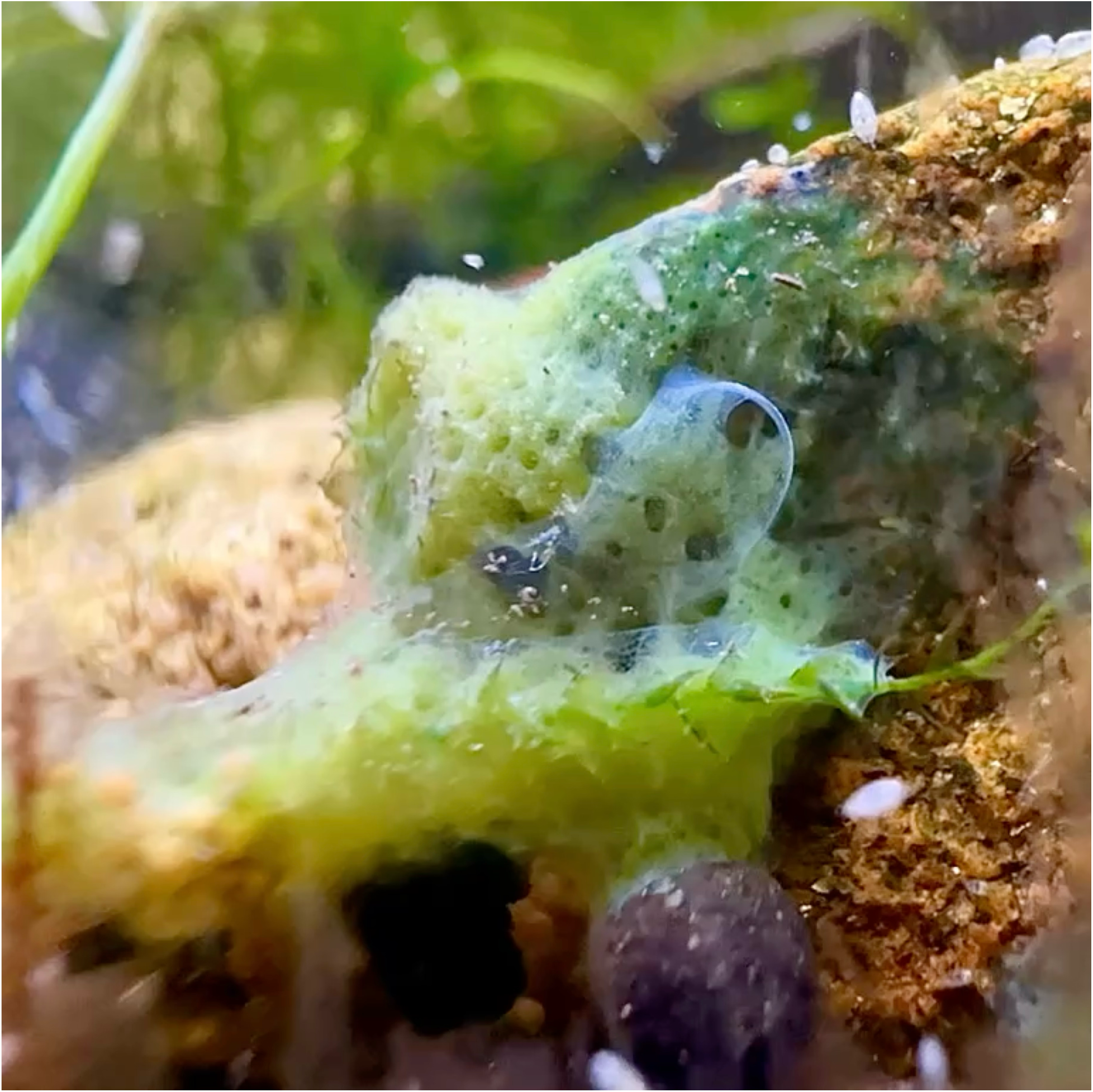
∼3-hour timelapse of aquarium-reared *E. muelleri.* Timelapse video (∼3 hours) of a ∼2-month-old, aquarium-reared *E. muelleri* individual grown from bulk sponge “wool” containing may individual gemmules. Timelapse highlights a wave of the “inflation-deflation” cycle of *E. muelleri*. This video additionally highlights waste excretion across the outer surface of the tent epithelium. An excurrent osculum is shown towards the top right of the individual in this video. Myriad macroinvertebrates (e.g., copepods, rotifers, limpets, etc.) are also present.

**Supplementary Video 2:**
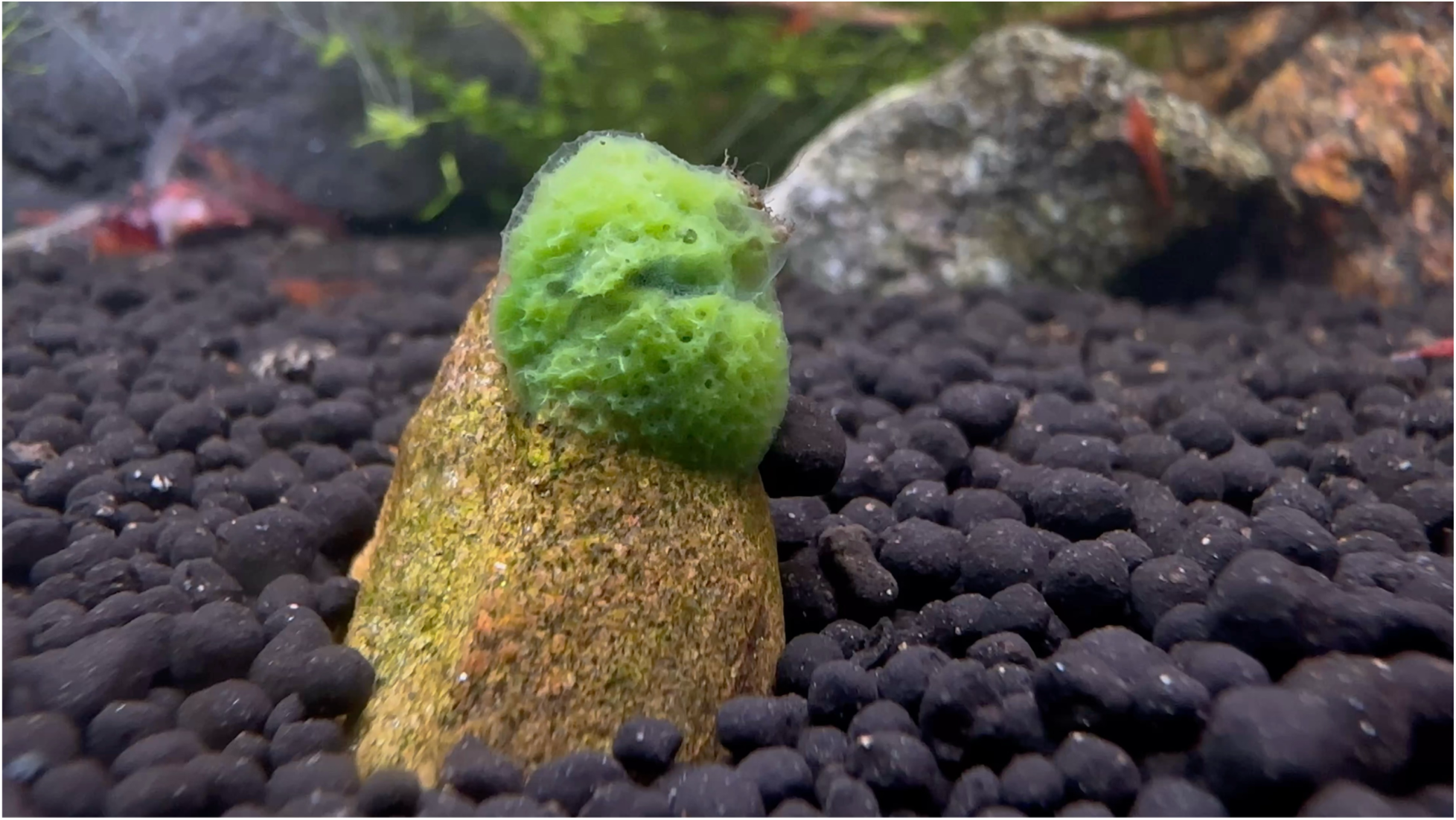
Real-time video of aquarium-reared *E. muelleri.* Real-time video (∼21 seconds) of a ∼5-month-old, aquarium-reared *E. muelleri* individual grown from bulk sponge “wool” containing many individual gemmules. The excurrent osculum is towards the top right in this video. Cherry shrimp (*Neocardinia* spp.) are seen readily throughout the aquarium.

## Notes

### Competing Interest Statement

The authors have declared no competing interest.

### Summary of Updates

Conducted new experiment to visualize bacteria on the surface of gemmules before and after sterilization treatments. We added more detailed replication descriptions. Added additional supplementary information to help orient the reader to the organization of the sponge body plan.

